# The application of Hadoop in Structural Bioinformatics

**DOI:** 10.1101/376467

**Authors:** Jamie Alnasir, Hugh P. Shanahan

**Affiliations:** Institute of Cancer Research, 123 Old Brompton Road, London, SW7 3RP, United Kingdom; Department of Computer Science, Royal Holloway, University of London, Egham, Surrey, TW20 0EX, United Kingdom

## Abstract

The paper reviews the use of the Hadoop platform in Structural Bioinformatics applications. Specifically, we review a number of implementations using Hadoop of high-throughput analyses, e.g. ligand-protein docking and structural alignment, and their scalability in comparison with other batch schedulers and MPI. We find that these deployments for the most part use known executables called from MapReduce rather than rewriting the algorithms. The scalability exhibits a variable behaviour in comparison with other batch schedulers, particularly as direct comparisons on the same platform are generally not available. We do note there is some evidence that MPI implementations scale better than Hadoop. A significant barrier to the use of the Hadoop ecosystem is the difficulty of the interface and configuration of a resource to use Hadoop. This will improve over time as interfaces to Hadoop e.g. Spark improve, usage of cloud platforms (e.g. Azure and AWS) increases and approaches such as the Workflow Definition Language are taken up.

## 1 Introduction

The Apache Hadoop project [73] is a software ecosystem i.e. a collection of interrelated, interacting projects forming a common technological platform [48] for analysing large data sets.

Hadoop presents three potential advantages for the analysis of large Biological data sets. In the first instance, it is designed for the analysis of large semi-structured data sets; secondly it is designed to be fault tolerant (in essence by ensuring a very large amount of overall redundancy) which becomes almost inevitable for sufficiently large numbers of processors; finally the MapReduce formalism for describing the data sets allows for the easy construction of work flows.

On the other hand, Hadoop also presents barriers to its adoption within the community for Bioinformatics and the analysis of structural data. In the first instance Hadoop runs as a series of Java libraries and hence there is a learning curve for any Bioinformatician or Structural Biologist who hasn’t used Java before, though we note that more recent additions to the Hadoop ecosystem such as Spark [67] have a wider range of languages. Correspondingly, unlike data parallel languages such as High Performance Fortran [77], Hadoop cannot be easily retro-fitted into a stable code base even if the original code is written in Java though it is possible to use Hadoop to deploy instances of executables in the same way that batch schedulers do. Finally, implementing Hadoop on a local cluster is not trivial and requires a significant level of expertise from the relevant systems administrator. As we note, this latter difficulty is obviated on cloud platforms such as Azure and AWS [66].

This paper reviews the range of work that has been done on Hadoop in Bioinformatics and in Structural Bioinformatics in particular. Specifically this paper will determine how stable these platforms are in comparison to other approaches such as batch schedulers and MPI (discussed in sections 1.3 and 1.4 respectively). The rest of this paper is organised as follows. In the first instance a brief overview of the Hadoop system as well as a description of batch schedulers and MPI. It then describes the Hadoop formalism. A brief review of the application of Hadoop in Bioinformatics is provided followed by an in-depth review of the application of Hadoop in Structural Bioinformatics. Finally this paper will draw some conclusions on the scalability of Hadoop and its application in Structural Bioinformatics.

### 1.1 Distributed computing architectures

### 1.2 Hadoop and MapReduce

Apache Hadoop is a software framework for distributed processing and storage which is typically installed on a Linux compute cluster to facilitate large scale distributed data analysis, though it can be run on a standalone single computer node usually for the purposes of prototyping. Hadoop clusters may be built using commodity hardware, for instance off the shelf equipment such as used in computer farms, and key features are **fault-tolerance** and **data-locality**. In the former case scaling up a cluster to add more machines and disks increases the probability of a failure occurring and hence systems must have a built-in redundancy to compensate for it. In the latter case data-locality provides the ability of the framework to execute code on the same node or at least the same rack of the cluster as where the input data resides. This reduces the amount of network traffic during processing thereby avoiding network bottlenecks [21]. The fault-tolerance and data-locality of Hadoop are made possible by its distributed file system (HDFS) [5] and by Hadoop’s resource scheduler YARN (Yet Another Resource Negotiator) [76] which is responsible for cluster management, in particular resource allocation and job scheduling.

The distributed data on HDFS is accessed programmatically using the MapReduce formalism, originally implemented in Java. In this formalism the distributed data accessed from HDFS is a set of *tuples* i.e. pairs of keys and values < *k_i_, υ_i_* >, 1 ≤ *i* ≤ *N* where *N* are the total number of data entries. For example, the entries in the Protein Data Bank (PDB) would be a set of pairs where *k_i_* would be a specific PDB id and *υ_i_* could be a single string with the corresponding PDB data in XML format. All operations are based on these tuples, either creating new tuples (i.e. through a *Map* step) or summarising the data stored in the tuples (i.e. through a *Reduce* step).

Extending the above example, via a *Map* step, a specific executable (e.g. a docking program run with a specific small molecule) could be run on each PDB entry to create a log file for each PDB entry. In the MapReduce formalism this means creating a new set if tuples < *k_i_,l_i_* > where *k_i_* is again the PDB entry and l_i_ is a single string with the log file. A *Reduce* step could then be applied on this second set of tuples to create a single tuple which carries some specific set of summary data (e.g. how many structures had a docking score greater than some threshold in the previous *Map* step).

The rise of the use of Hadoop has mirrored the increasing use of cloud platforms. MapReduce is offered as a Platform as a Service (PaaS) by all of the major cloud-service providers (Amazon AWS, Google Cloud and Microsoft Azure) [20].

### 1.3 Batch schedulers

A batch-scheduler (also referred to as a job-scheduler or workload management software), is a central software component of a compute cluster that manages the workload of computational jobs on a cluster and allocates cluster resources to them [30]. We will refer to the technique as batch-scheduling and the central software component as the job-scheduler. Generally a computational job on a batch-scheduled system is a normal user program that runs on a single compute node, but can also be a more specialist distributed program that comprise of components written to run on multiple nodes which communicate by passing messages, for example using message passing interface (MPI) (discussed in the following section). Computational jobs are submitted in a batch to the job scheduler which adds them to a queue. The job scheduler decides on how to prioritise competing jobs, what resources to allocate to the jobs. The jobs are then submitted by the job scheduler to compute nodes of the cluster using a scheduling algorithm [12].

Batch-scheduled cluster systems couple job flow control with the ability to reserve (and limit) the allocation of cluster resources such as, for example, CPU cores, physical RAM, virtual memory and execution time. However batch-scheduled clusters offer only “course granularity” control of concurrency at the job-level (unlike MPI systems) and does not render the same level of fault-tolerance and data-locality through lack of a distributed file system such as HDFS that Hadoop provides (discussed above). Batch-scheduled cluster systems use the job scheduler to deploy “whole” executable programs to compute nodes of the cluster which may or may not run in a parallel fashion - for instance a single program when submitted will run on only one node, while multiple submitted jobs may run either on a single node or be distributed across multiple nodes depending on the load on each compute node and the scheduling rules set.

### 1.4 MPI

High performance computing (HPC) systems are typically reliant on a high degree of inter-process communication. The Message Passing Interface (MPI) is a standard for the development of parallel programming software and libraries for parallel computing architectures that standardises syntax [47]. MPI facilitates concurrent programming by specifically dictating the standard syntax to be used for the messages passed between communicating processes. MPI supports a wide variety of architectures such as multiple computers with distributed memory, shared memory multiple processors and heterogenous combinations of these.

MPI offers an extremely fine granularity of control over the processes involved in the execution of parallel, concurrent programs running across networked computers in clusters. Although MPI is extensively used in high performance computing, it can be used on clusters of standard machines or workstations. MPI requires the programmer to explicitly handle parallel functionality at a lower level than for instance Hadoop which automates parallelism of users programs through the MapReduce formalism. Whilst MapReduce has parallels to MPI programming, especially in relation to MPI functions *scatter* and *reduce,* it offers automatic parallelism, as well as data-locality and fault-tolerance (discussed previously) [27].

## 2 Applications of Hadoop in Bioinformatics

The emergence of MapReduce based platforms that we have discussed such as Hadoop and Spark have not been overlooked by researchers in different areas of bioinformatics. A number of projects within the Apache Hadoop ecosystem find useful application in bioinformatics [73]. These include the data-warehousing framework Hive [74] which has an SQL type query language, the high level data-flow language Pig [55] which compiles scripts into sequences of MapReduce steps for execution on Hadoop, the machine-learning and clustering facilities offered by Mahout [41], and HBase a distributed scalable database [18]. All of these projects utilise Hadoop’s cluster infrastructure and distributed file system and therefore gain from the scalability and fault-tolerance inherent in their design, as discussed earlier.

In terms of software applications MapReduce has been employed for a variety of problems in processing biological and sequencing datasets. Some notable projects in the area of sequence alignment are, Cloudburst [63] and CloudAligner [52], which are both based on the RMAP alignment algorithm [70], and CloudBlast [43] which is based on the BLAST algorithm [3]. It is noteworthy that MapReduce can be especially suited for, for example the construction of a de Bruijn graph for de *novo* genome assembly. For example, Contrail is able to build adjacency lists for all the *k-mers* in the genomic sequence reads and then uses distributed aggregation functions such as *reduce* to compress simple chains of length N in *O*(*log*(*N*)) rounds using a randomized parallel list ranking algorithm [64].

There are also tools implemented in MapReduce for the analysis of assembled sequencing data, for instance Crossbow [36] is designed for SNP (Single Nucleotide Polymorphism) detection. It uses the Bowtie [75] and the SNP caller SOAPsnp [38]. Differential expression (using RNA-Seq) can be measured using the Myrna software pipeline [35] - pipelines are data-flows comprising of sequential steps in which bioinformatics software are applied to the data [37].

Additionally, a number of programming libraries that facilitate the manipulation and processing of sequencing data file formats such as SAM Sequence Alignment Map and BAM (Binary Alignment Map) have arisen such as the Java based libraries Genome Analysis Toolkit (GATK) [45] developed by the Broad Institute and Hadoop-BAM [53] as well as the Scala based SparkSeq [80] (discussed below). The GATK provides functions for data management in the form of data access patterns, namely the low level implementation is separated from higher level functions, and also provides functions for analysis calculations. The Broad Institute have also developed a Workflow Definition Language (WDL) for use in data analysis pipelines (discussed in the next section). It is a high-level language that is designed to be human readable and writable. It allows researchers to describe analysis tasks, daisy-chain tasks into workflows, and utilise advanced features such as parallelization [8]. WDL was developed out of the necessity for standardisation amongst a number of different pipeline solutions, thereby providing a universal standard. In order to execute analysis pipelines written in WDL, an execution engine is necessary. Cromwell is such an engine, also designed by the developers of WDL, to run on many platforms (Locally, HPC or Google - support for other platforms such as Microsoft Azure and AWS is forthcoming) and can scale elastically to workflow needs [7].

The provision of pipeline development specifically for the Hadoop platform is also available. For instance, SparkSeq is a MapReduce library for building genomic analysis pipelines using Scala on Apache Spark. Whilst Scala is supported on the Spark platform it lacks the same user base in bioinformatics as it enjoys amongst the data analytics and machine learning communities. Given the vast amounts of sequencing data being produced [72, 79], the purpose of these tools is to exploit the scalability that is characteristic of MapReduce and which the Hadoop and Spark platforms offer, and this offsets any difficulty in developing or re-writing applications using the MapReduce paradigm. However, the development of universal standards, such as WDL offers researchers a means of utilising tools developed for such platforms in a more user-friendly way.

## 3 Applications in Structural Bioinformatics

The Protein Data Bank (PDB) is an archive of data describing the 3D shapes of proteins, nucleic acids, and molecular-complex assemblies derived from x-ray crystallographic, Nuclear Magnetic Resonance spectroscopy (NMR) and electron microscopy techniques [1, 6]. It also serves as a portal for structural genomics [34].

There are also a number of applications for Structural Bioinformatics implemented using MapReduce on Hadoop, specificially to carry out high-throughput analyses of such data sets which will be discussed. Whilst the focus of this section will be on systems developed for the Hadoop platform, for purposes of comparison, it will also refer to similar systems implemented on other platforms.

### 3.1 Molecular docking

Molecular docking typically involves simulating the electrostatic interactions between a ligand (often a potential drug molecule) and a target protein [49, 46]. It is used to score ligands on their affinity to the target, usually for the purposes of drug development - a process that is complex, time-consuming and expensive [50, 61].

#### 3.1.1 Docking of protein-ligand complexes on Hadoop

A number of molecular docking applications have been implemented to exploit the Hadoop platform. For example, [15] at the Oak Ridge National Laboratory in the US, have utilised AutoDock4 on a private 68 node Hadoop cluster to perform the docking of 2,637 compounds from the Directory of Useful Decoys (DUD) database [24], against the Human estrogen receptor alpha agonist protein (PDB entry 1L2I, [68]). They used the DUD database because it contains ligands that bind to the target, as well as chemically similar ligands that do not (decoys). This allowed them to test the reproducibility of the docking experiments - they found that the results of running AutoDock on Hadoop were consistent with the experiments of [24], specifically that the percentage of known binding ligands correlated with the percentage ranked in the DUD database. In their configuration they used 10 mappers per node on the 57 nodes of their cluster that were dedicated to run MapReduce Tasks giving 570 mappers running in parallel. This resulted in a 450x speed-up of AutoDock in performing the docking task on Hadoop as compared with utilising AutoDock itself to manage the parallelisation. Furthermore, they report that 95.59% of CPU time is used by AutoDock, and, therefore, there is less than a 5% overhead in running AutoDock in a Hadoop map process, and that, as the tails seen in the graphs of the CPU load were steep, this indicates that job initialisation and termination are not resource intensive.

As a comparison, [83] conducted the same experiment using the DUD database with MPI and a multi-threading parallel scheme at an extremely large scale (15,408 CPUs). They found that VinaLC scaled very well up to with an overhead of only 3.94%. 17 million flexible compound docking calculations were completed on 15,408 CPUs within 24 hours. 70% of the targets in the DUD data set were recovered using VinaLC. These applications, and the others we will discuss are listed in Table ?? for comparison.

Another system for protein-ligand binding on Hadoop, developed by [25] is a scalable docking service called Cloud-PLBS (Cloud Protein Ligand Binding Service), which utilises the SMAP docking tool [81]. Their system employs an additional virtualisation system, whereby the Hadoop slave nodes run on virtualised machines and are instantiated depending on the input job requirements. In terms of benchmarking performance, they compared stand-alone, sequential processing of protein-ligand pairs using SMAP, with parallel execution of SMAP within a Hadoop map function - specifically 2, 4, 6 and 8 mappers. They observed that in docking 40 protein-ligand pairs, reduction in execution time using Hadoop vs. stand-alone for 2, 4, 6 and 8 mappers was 33.92%, 56.97%, 70.21%, 77.65%, respectively.

They also tested the fault-tolerance of Hadoop in running their protein-ligand pair docking system by simulating task failures in 50% of the map steps by removing node(s) from service.

They observed that the docking jobs still completed. As discussed previously, fault-tolerant distributed computation is a feature of Hadoop based applications, and this resilience in the execution of tasks is important, because the likelihood of a node failing increases with the scaling-up of a cluster. Fault-tolerance is also desirable in web-based services such as Cloud-PLBS, which serve to automate computational jobs and present the results to the user, without requiring third-party intervention to rectify failed jobs. However, it should be noted that no reference to source code for their system is provided in their paper, and the Cloud-PLBS service at is no longer available.

#### 3.1.2 Clustering of protein-ligand complexes

One of the challenges in the field of molecular docking studies arises from the requirement to search the conformational space of protein-ligand complexes generated across docking experiments. This is necessary to select the most likely conformations of protein-ligand complexes, and, therefore, the putative ligands (potential drug molecules) which partake in these interactions. In such experiments large numbers of protein-ligand complexes are generated, docked, and scored [16], and it is, therefore, necessary to select a subset of putative ligands based on significant protein-ligand interactions.

Estrada *et al.* observe that selecting the native conformation, based on the assumption that the lowest energetically scored conformation (as computed by an energy function) represents the native binding of the ligand and protein, is not reliable, even in larger sets of conformations. This is often due to non-native ligand-protein complexes generating falsely low energy scores. They point out that, whilst hierarchical clustering techniques are a logical way of addressing this problem - as the lowest scoring, most densely populated clusters overlap with native conformation - most clustering algorithms are computationally expensive, and scale poorly with large datasets. They implemented a system using MapReduce on Hadoop to address this issue. They used two datasets, of size 5 TB (3,872 million ligand conformations) and 1 TB (768 million ligand conformations), generated from the Docking at Home volunteer grid computing project (Docking@Home) [17], which utilised CHARMM [9].

The examples discussed in the section 3.1.1 did not fully implement their solutions in MapReduce. This would have involved implementing (or re-implementing) algorithms using MapReduce, but instead exploited Hadoop’s *map* step to encapsulate and execute external applications. The method discussed in [16] however, is fully implemented in MapReduce.

A map step is used which geometrically reduces the conformational space. This is stored in an Octree data structure [62], together with a unique identifier (an Octkey) used for traversal. This is achieved by projecting the *x, y, z* components of the conformations onto a 2D plane, and calculating their gradients (for *x*,*y*, and *z*) which are then encoded into a single point in the Octree. A reduce step is used to aggregate conformations in the Octree. Further MapReduce operations are then used to traverse the Octree using the Octkey identifier.

In order to compare the accuracy of their Hadoop-based Octree method (for selecting native conformations from the ensemble of complexes) against other approaches, namely Hierarchical clustering and Minimum Energy selection, they docked 100,000 protein-ligand complexes each for HIV, Trypsin, and P38alpha. They obtained 80%, 75% and 25% accuracy for Hadoop based Octree, Hierarchical Clustering and Minimum Energy methods, respectively.

They also examined the accuracy of selecting native conformations from the cross-docking data in the Docking@Home datasets, whereby each conformation of the ligand in the set of complexes is docked with each conformation of protein. In doing so, they compared their Octree method with the Energy Minimum approach, and observed 43.8% and 5.8% accuracy, respectively.

In testing the scalability in processing the 5 TB dataset which, as discussed contains 3,872 million ligand conformations, they used a Hadoop cluster where each node possesses 32 cores (4x Octacore AMD Opteron 2.4 GHz), and up to a maximum of 32 nodes were available. The range of the scaling used was 1 node of 32 cores (to process 121 million conformations) to 32 nodes with a total of 1024 cores (to process the full dataset) and analysed 1, 2, 4, 8, 16, and 32 nodes - in all cases, the number of ligand conformations processed per core was 3.8 million.

It is not stated in their paper how many map processes were running per core, but it is assumed that it is 1 map step per core. Whilst they demonstrated their method was amenable to scaling, they observed an appreciable decrease in parallel efficiency with the increase in cores, from 99.1% down to 43.8% for 64 cores (2 nodes) and 1024 (32 nodes), respectively. This appears to be due to the increased overhead in communications between the processes as the number of processes increases (communications to computation ratio).

A similar application [58] was developed on the Hadoop platform that partitions the results of molecular dynamics simulations. The trajectories of atom positions, velocities and energies as a function of time are clustered, as large datasets. This method yields important information about the most probable conformations of proteins in ensembles. Their system employs the GROMOS algorithm [65], which is not inherently parallel, by implementing it as a series of map and reduce functions so as to utilise Hadoop. They tested their parallelised MapReduce implementation of the GROMOS algorithm on a Hadoop cluster comprising of 1 master and 3 slave nodes, each comprising two hexa-core Xeon E5645 CPUs 32 GB of RAM and 2 TB of disk space. They observed up to 10 and 7 times speed-up (over using sequential GROMOS) of the first and second phases, and final two phases of their algorithm, respectively.

A docking application also relevant to our discussion, although not implemented in Hadoop, has been developed by [54] using a scientific workflow management tool, SciCumulus deployed on AWS. Their system employed molecular docking, using AutoDock4 and Vina, on their platform to explore both drug discovery and scalability. Their drug discovery objective was the identification of putative drug ligands that bind to Cysteine Proteases of Protozoan genomes utilising 10,000 protein-ligand complexes. This aims to facilitate the development of drugs for the Neglected Tropical Diseases (NTDs). In investigating the scaling-up of the computational task, they utilised up to 32 heterogeneous nodes (containing varied numbers of cores) to include a total of 128 Amazon AWS EC2 cores. They observed an almost linear relationship between number of nodes and speed, but this plateaued as they approached the maximum of 32 nodes, suggesting less benefit in scaling beyond this. They point out that this is likely due to more complicated load balancing in the set of heterogeneous nodes. The result of their docking experiments using the Cysteine Protease-ligand complexes identified 287 and 355 putative ligands for AutoDock4 and Vina, respectively. It is important to note, however, that these potential drug molecules have, on average, RMSDs greater than 2 — 3 Å (Angstroms) which is the maximum accepted value for a useful result.

### 3.2 Structural Alignment

The alignment of proteins by structure, as opposed to by sequence, is a computational technique used to identify homologous polymer structures within proteins that may be conserved between proteins. The technique facilitates the study of the structural and evolutionary relationships of proteins with low sequence similarity [19, 56]. A variety of of algorithms have been developed to perform structural alignment of proteins, such as, for example, STRUCTAL (Structural Analysis Algorithm), DALI (Distant Alignment) [22], CE (Combinatorial Extension) [69], VAST (Vector Alignment Search Tool) [19], and FATCAT (Flexible structure Alignment by Chaining Aligned fragment pairs allowing Twists) [82], SSAP (Sequential Structure Alignment Program) [57], and MUSTANG (Multiple Sequence Alignment Algorithm) [32]. The technique has been applied to the study of protein binding sites, and solvent exposed surfaces (these effect the energetics of protein-ligand conformations) [42, 33, 39].

Structural alignment algorithms are usually computationally complex, [31] present a method which runs at best in approximately Polynomial time, but they also point out that approximations are often used. There is also a need to apply such techniques at scale. For example, Hadoop has been employed by [39] who implemented structural alignment for binding site prediction. Using a test sets of 200 and 48 ligand-protein complexes, they were able to achieve 93% and 98% accuracy, respectively, and were able to improve the efficiency of the experiment by exploiting parallelisation. A service for structurally aligning pairs of proteins has also been implemented for the Hadoop platform by the developers of Cloud-PLBS [26] (discussed in the previous section 3.1). It utilises the same distributed architecture as Cloud-PLBS, that is, individual Hadoop nodes running within their own VM, and each VM running a *map* and *reduce* process. As with Cloud-PBS, a web-interface is used to enable the user to provide input of two PDB files by their PDB-ID. To perform the structural alignment they state their system uses the DALI and VAST algorithms. Whilst the authors do detail the basis of RMSD (Root Mean Square Deviation) in structural alignment algorithms, and discuss refinement methods in their paper, they claim to implement these algorithms in MapReduce. As previously noted with Cloud-PLBS, there is no source code available, and the corresponding web service is unavailable. It is highly likely that, given the complexity of these algorithms and that there are already implementations available, the same method used in Cloud-PLBS - execution of an external program within a map step - is employed.

A similar bioinformatics SaaS (Software-as-a-Service) for structural alignment of proteins, has been developed for the Microsoft Azure platform - Cloud4Psi developed by [51]. Their service utilises three newer algorithms that are implemented in the BioJava project, and which are derived from CE (jCE), and FATCAT (jFATCAT-rigid and jFATCAT-flexible) [60]. They tested their system on a subset of 1,000 PDB structures for both scalability and reproducibility. For scalability, two different scaling methods were compared: *horizontal-scaling* (i.e. by adding more nodes to the system) and *vertical-scaling* (i.e. by using nodes with more CPU cores). They found that, whilst both scaling methods increased the n-fold speed-up for each of the three algorithms, both suffered a decrease in the performance gains - for horizontal scaling, this was found to be the result of increased disk I/O due to multiple nodes utilising a shared VHD (Virtual Hard-Disk), and for vertical scaling this was due to an increase in processes on the same node (due to higher specification of each node), resulting in increased CPU utilisation. As the horizontal scaling method suffered less from this effect, it was the method chosen. For reproducibility of results, they found that each of the three algorithms produced the same results independent of cluster configuration and scaling used.

### 3.3 Other Structural Bioinformatics applications using Hadoop

Large-scale processing of molecular data is desirable in both applications. Such techniques facilitate the *in-silico* study of vast arrays of molecular compounds and macromolecular structures that are available from large molecular databases, which are also increasing in size and diversity [14, 59, 2, 78, 1, 6]. A number of scalable Structural Bioinformatics methods have been provided by the bioinformatics Group at UCL (University College London) as web-based services through their Protein Analysis Workbench [11]. These are accessible via SOAP (Simple Object Access Protocol), and XML-RPC (Extensible Markup Language-Remote Procedure Call) protocols. Importantly, the most commonly used methods have also been deployed as Java packages specifically for the Hadoop platform. This includes PSIPRED for protein structure prediction [44], GenTHREADER for protein fold recognition method using genomic sequences [28], BioSerf - a homology modelling protocol, MEMSAT for improving accuracy of transmembrane protein topology prediction [29], DomPred for protein domain boundary prediction [10], MetSite for predicting clusters of metal-binding residues [71], and FFPred which uses a machine learning approach for predicting protein function [40].

## 4 Conclusions

This purpose of this review is to give an insight into the impact that Hadoop and the MapReduce formalism has in Structural Bioinformatics. This is a apposite moment to consider this as there have been a number of different applications of Hadoop in the field and Hadoop (and the wide variety of different applications that have been built on it) has become more stable and accessible.

As noted previously the adoption of Hadoop is not a trivial step, particularly for a Structural Bioinformatics lab that already has extensive experience in using traditional batch schedulers running on a local cluster. Rewriting stable codes to make them use the MapReduce formalism at the lowest level would require substantial effort though using MapReduce to call executables requires much less effort and can still make use of the fault-tolerance and data locality features of Hadoop.

This article has focussed on

- determining the breadth of cases where Hadoop has been used,
- how well it scales and
- how dependent the installation of Hadoop is on its configuration (which is indicative of the difficulty one would have in installing Hadoop).

In the first instance we see a range of traditional high-throughput applications in Structural Bioinformatics (e.g. docking and structural alignment) where Hadoop has been employed.

With respect to scaling, the publications reviewed here indicate that some adjustment of parameters have been made, but these largely focused on how the applications scale with the number of nodes. They show that performance is linear though the gains in performance tend to reduce as the systems are scaled up. In molecular docking, [16] observed a fall in parallel efficiency of 55.3% (99.1% - 43.8%) when scaling from 2 nodes to 32 nodes. In structural alignment, [39] observed that performance degraded slightly after 8 mappers was increased to 30. Furthermore, this trend has also been observed on the MS Azure platform we have discussed for comparison - in scaling Cloud4Psi, also a structural alignment application [51], observed that horizontal scaling resulted in performance degradation as a result of an increase in nodes sharing a virtual disk, and that vertical scaling resulted in performance degradation as a result of increased CPU utilisation (due to more processes running per node).

The platform, and method of distribution, also dictate performance and scalability. In observing two identical docking operations on the DUD database, one using a 1,088 core Hadoop cluster [15], and the other using 15,408 cores with a mixed parallel MPI implementation of AutoDock [83], the Hadoop cluster took 69 hours, and the MPI implementation completed within 24 hours. Whilst this is certainly a result of the number of cores, the MPI system scaled better, with only very slight degradation in performance after 6,000 CPU cores. Although this comparison involved the same docking operation and dataset using different platforms, currently, there are no comparisons of performance between Hadoop and batch-schedulers on the same cluster apparatus in the literature.

The comparison of vertical and horizontal scaling in Cloud4Psi indicates significant change in performance, so configuration *is* important. As noted with the Cloud4Psi example, there can be a significant variation depending on configuration. Performance gains across applications, therefore, are dependent on configuration.

As Hadoop platforms stabilise, the significant advantage of its employment is of using a platform where the computation is expressed explicitly in terms of an algebra. This makes building workflows easier by allowing the developer to concentrate on the calculation, rather than the process such as WDL [8]. Nonetheless, there is a significant gap in take up as such systems remain difficult to deploy.

The Hadoop ecosystem is rapidly evolving (as we have seen with the introduction of Spark) and such gaps will reduce over time. On the other hand, it will also require regular release upgrades. These can be potentially be difficult to deploy, and often add new components to the ecosystem which increase the potential to introduce bugs that may affect different areas of the system.

To address this, organisations such as Cloudera and Hortonworks [13, 23] provide supported Hadoop-stack releases, and cloud-service providers such as Amazon offer managed-services, for example Elastic Map Reduce (EMR) [4]. Implementing systems on the Hadoop platform, as discussed, also requires specialist programming knowledge of MapReduce, and if the relevant Hadoop cluster is maintained in-house, specialist skills in maintaining an Hadoop cluster are also required. For this reason, managed services are often utilised by enterprise companies because such systems have been deployed and tested by technical experts and therefore mitigates risks, and dispenses the need to employ or train in-house skilled personnel to maintain a Hadoop cluster.

## Acknowledgements

The research in this article was made possible from support from the Department of Computer Science, Royal Holloway, University of London.

